# Back pain, mental health and substance use are associated in adolescents

**DOI:** 10.1101/274605

**Authors:** SJ Kamper, ZA Michaleff, P Campbell, KM Dunn, TP Yamato, RK Hodder, J Wiggers, CM Williams

## Abstract

**Background:** During adolescence, prevalence of pain and health risk factors such as smoking, alcohol use, and poor mental health rise sharply. While these risk factors and mental health are accepted public health concerns, the same is not true for pain. The aim of this study was to describe the relationship between back pain and health risk factors in adolescents.

**Methods:** Cross-sectional data from the Healthy Schools Healthy Futures study, and the Australian Child Wellbeing Project was used. The mean age of participants was 14-15 years. Children were stratified according to the frequency they experienced back pain over the past 6 months. Within each strata, the proportion of children that reported drinking alcohol or smoking in the past month and the proportion that experienced feelings of anxiety or depression was reported. Test-for-trend analyses assessed whether increasing frequency of pain was associated with health risk factors.

**Results:** Data from approximately 2,500 and 3,900 children in the two studies was analysed. Larger proportions of children smoked or drank alcohol within each strata of increasing pain frequency. The trend with report of anxiety and depression was less clear, although there was a marked difference between the children that reported pain ‘rarely or never’, and those that experienced back pain more frequently.

**Conclusion:** Two large, independent samples show Australian adolescents that experience back pain more frequently are also more likely to smoke, drink alcohol and report feelings of anxiety and depression. Pain appears to be part of the picture of general health risk in adolescents.

**What is already known on this subject?:** The prevalence of back pain rises steeply during the adolescent years, and is responsible for considerable personal impact in a substantial minority. During this time, indicators of adverse health risk such as smoking, alcohol use, anxiety and depression also increase in prevalence. Pain and lifestyle-related health risk factors can have ongoing consequences that stretch into adulthood.

**What this study adds?:** This study shows a close relationship between increasing pain frequency, and tendency to engage in health risk behaviours and experience indicators of poor mental health in adolescents. This study shows that pain may be an important consideration in understanding the general health, and health risk in adolescents.

## Background

Substance use and psychological distress are causes of concern in adolescents. Population-based surveys report substantial prevalence of alcohol, tobacco and illicit substance use in teenagers in numerous countries including the USA,^25^ the UK,^16, 30^ and Australia.^5^ This is of concern because the developing brain may be acutely susceptible to negative influences of toxic substances, and use in early adolescence may increase the risk of substance use disorders and mental health problems in later life.^9, 31^ Psychological distress, including anxiety, stress and depression are major contributors to the burden of disease among adolescents. Major depression, anxiety, and conduct disorders rank 3^rd^, 4^th^ and 7^th^ of all conditions for years lived with disability in 15-19 year-olds.^18^ Similar to substance use, these conditions are not only responsible for a concurrent burden, but are drivers of poor health into adulthood.^4^ It is accepted that substance use issues and mental health are closely related, although causal associations are undoubtedly complex.^1^

During adolescence, the prevalence of musculoskeletal (MSK) pain (pain arising from the bones, joints or muscles) in general, and back pain in particular rises steeply.^21, 22^ Although often dismissed as trivial and transient, there is mounting evidence that adolescent back pain is prevalent,^6, 29^ associated with a large disability burden in its own right,^18^ and responsible for substantial health care utilisation,^12^ medication usage,^8^ school absence,^27^ and interference with physical and day-to-day activities.^22^ Of further concern is the fact that persistent episodes of pain during adolescence appear to be related to development of chronic pain in adulthood.^2, 10, 14^

Previous studies have reported associations between MSK pain, substance use, and poor mental health, but the relationships between them are unclear. Evidence from a systematic review shows a small but robust relationship between smoking and chronic back pain among teenagers.^28^ Studies investigating the relationship between pain in adolescents and alcohol consumption report inconsistent results including; positive associations,^11, 13^ no association,^23^ and a study that reported adolescents with pain to be less likely to drink alcohol.^24^ There is robust evidence that demonstrates links between MSK pain and indicators of poor mental health,^17^ for example; between recurrent MSK pain and anxiety,^3^ MSK pain and internalizing (anxiety, depression, withdrawal) and externalizing (aggression, rule-breaking) behaviours,^20^ as well as evidence of mental health factors associated with, and influencing substance use. Greater clarity is required on the relationships between MSK pain, substance use, and poor mental health, specifically regarding pain type, for example transient pain compared to pain that is persistent.

Much of the epidemiological research into MSK pain in children does not distinguish between the presence of ‘any’ pain, and pain that persists or is experienced frequently. This may contribute to inconsistency in the body of literature. Because pain is a part of everyday life, for children as with adults, it is important to distinguish between pain of consequence (e.g. persistent pain, frequent pain, or pain associated with activity limitation, school absenteeism, health care seeking) and trivial, transient episodes. Stratifying pain by frequency offers one way of attempting to make this distinction.

The aim of this study was to determine whether adolescents that experience back pain more frequently were also more likely to report other indicators of adverse health risk e.g. alcohol use, smoking, school absenteeism and indicators of poor mental health e.g. depression and anxiety.

## Methods

We used data from two samples of adolescents to address the study aim. We chose to examine the study aim independently in the two datasets as a check on generalisability of the findings.

The Healthy Schools Health Futures study (HSHF) is a cluster randomised controlled trial testing the effectiveness of a resilience intervention on substance use in high school children. The HSHF trial included 32 high schools in the Hunter New England region of NSW, Australia. Schools were eligible if they were: 1) located in disadvantaged Local Government Areas (score of less than 1000 on the SEIFA Index of Socio-Economic Advantage/Disadvantage); and 2) co-educational with at least 400 students in Years 7 to 10. Comprehensive details of the study methods have been reported elsewhere.^15^ At follow up for the HSHF trial in 2015, Year 9 students provided data for the current study. Based on cultural advice processes within the project governance, including consultation with a cultural advice group, students identifying as Aboriginal and/or Torres Strait Islander were not asked to provide information on pain outcomes. This was because Aboriginal students were already receiving additional items relating to community and culture, and further addition to the survey was not reasonable. Surveys were administered online during class time.

Ethical approval for the original study was obtained from the Hunter New England Health Human Research Ethics Committee (Ref no. 09/11/18/4.01), The University of Newcastle Human Research Ethics Committee (Ref no. H-2010-0029), the Aboriginal Health and Medical Research Council (Ref no. 776/11), the New South Wales Department of Education and Training State Education Research Approval Process (Ref no. 2008118), and the relevant Catholic Schools Offices. The trial is registered with the Australia and New Zealand Clinical Trials Register (Ref no. ACTRN12611000606987).

The Australian Child Wellbeing Project (ACWP) is a cross-sectional survey of Australian schoolchildren. The ACWP is a nationally representative survey of children in years 4, 6 and 8 conducted in 2014. Data from children in year 8 attending 180 government, catholic and independent schools across all states in Australia were used for this study. Surveys were administered online, with audio support for children with low literacy levels. Comprehensive details of the study methods are available elsewhere.^26^

Ethical approval for the ACWP project was obtained from the Australian Council for Educational Research, Flinders University and the University of NSW. Further permissions were received from state/territorial authorities and dioceses and participation required written informed consent from parents and children.

### Measures

Back pain was measured slightly differently, but comparably in the two datasets. The questions were as follows:

- HSHF: “In the last 6 months, how often have you had any pain in your back?” (never, rarely-once or twice, about every month, about every week, more than 1x/week, about every day)
- ACWP: “In the last 6 months, how often have you had backache?” (rarely or never, about every month, about every week, more than 1x/week, about every day)

Adverse health risk indicators were measured as per below, creation of dichotomous indicators was defined prior to analysis:

- Smoking:
  - HSHF “Have you smoked a cigarette in the last 4 weeks?” (no, yes).
  - ACWP “On how many occasions have you smoked in the last 30 days?” dichotomised: never // 1-2 times, 3-5, 6-9, 10-19, 20-39, 40 or more.
- Drinking:
  - HSHF “In the last 4 weeks, how many times have you had 5 or more alcoholic drinks in row?” dichotomised: none // once, twice, 3-6 times, 7 or more times.
  - ACWP “On how many occasions have you been drunk in the last 30 days?” dichotomised: never // 1-2 times, 3-5, 6-9, 10-19, 20-39, 40 or more.
- Depression:
  - HSHF “How much of the time during have you felt down hearted or blue?” dichotomised: none // a little bit of the time, some of the time, a good bit of the time, most of the time, all of the time.
  - ACWP “In the past 6 months how often have you (had the following) feeling low?” dichotomised: rarely or never // about every month, about every week, more than 1x/week, about every day.
- Anxiety:
  - HSHF “How much of the time during the past month have you been a very nervous person?” dichotomised: none // a little bit of the time, some of the time, a good bit of the time, most of the time, all of the time.
  - ACWP “In the past 6 months how often have you (had the following) feeling nervous?” dichotomised: rarely or never // about every month, about every week, more than 1x/week, about every day.
- Missed school:
  - ACWP “Last term how many times have you missed school?” dichotomised: never, hardly ever // about 1x/week, most days, every day.

We reported age, and selected sociodemographic measures to characterise the sample.

### Analysis

Data were analysed using the Mantel-Haenszel test-for-trend using SPSS version 24. The analysis provides an indication of whether the odds of the adverse health indicators rise incrementally with increasing frequency of pain. Analyses were conducted separately in each dataset, and the findings compared to provide an indication of generalisability.

## Results

### Sample characteristics

Characteristics of the participants are presented in Table 1, both samples are similar with respect to age, gender proportion, and language spoken at home. The fact that the HSHF study preferentially recruited schools from lower socioeconomic areas is reflected in the distribution of socioeconomic status variable, frequency of back pain was also higher in this dataset.

**Table 1.** Participant characteristics

|  | **ACWP** | **HSHF** |
| --- | --- | --- |
| **Age range** | 14-15 years<br>n=3896 | 14-16 years<br>n=2492 |
| **Female gender (%)** | 48.8%<br>n=3896 | 49.3%<br>n=2506 |
| **Socioeconomic status (%)** | 18.0% low<br>36.1% middle<br>45.9% high<br>n=3896 | Quintile 1; 12.5%<br>Quintile 2; 30.9%<br>Quintile 3; 45.6%<br>Quintile 4; 10.5%<br>Quintile 5; 0.4%<br>n= 2506 |
| **English spoken at home (%)** | 92.9%<br>n=3893 | 93.0%<br>n=2503 |
| **Quality of life\* (mean (SD))** | 7.62 (1.69)<br>n=3881 | - |
| **Overall subjective health (%)** | 34.8% excellent<br>54.9% good<br>9.0% fair<br>1.4% poor<br>n=3874 | - |
| **Back pain frequency** |  |  |
| Rarely or never | 51.9% | 66.7% |
| Every month | 22.5% | 8.0% |
| Every week | 11.1% | 8.7% |
| >1x / week | 8.0% | 9.0% |
| Every day | 6.5% | 7.7% |
|  | n=3608 | n=2075 |

**ACWP:** Australian Child Wellbeing Project; **HSHF:** Healthy Schools Healthy Futures study; n= total number of children who responded to the survey item;
\* Quality of life: 0=worst; 10=best

### Primary analysis

In both the samples, the proportion of participants reporting smoking, drinking and missing school (ACWP data only) rose incrementally with increasing frequency of pain. The test-for-trend was significant for all risk factors in both samples. With respect to the mental health indicators, the proportion reporting feelings of depression or anxiety was much lower in the group that never or rarely experienced pain, but did not increase substantially in the other groups. Test-for-trend analyses for these mental health factors were statistically significant (Table 2, Figure 1). The absolute rates for all health risks were generally higher in the HSHF sample.

**Figure 1.**
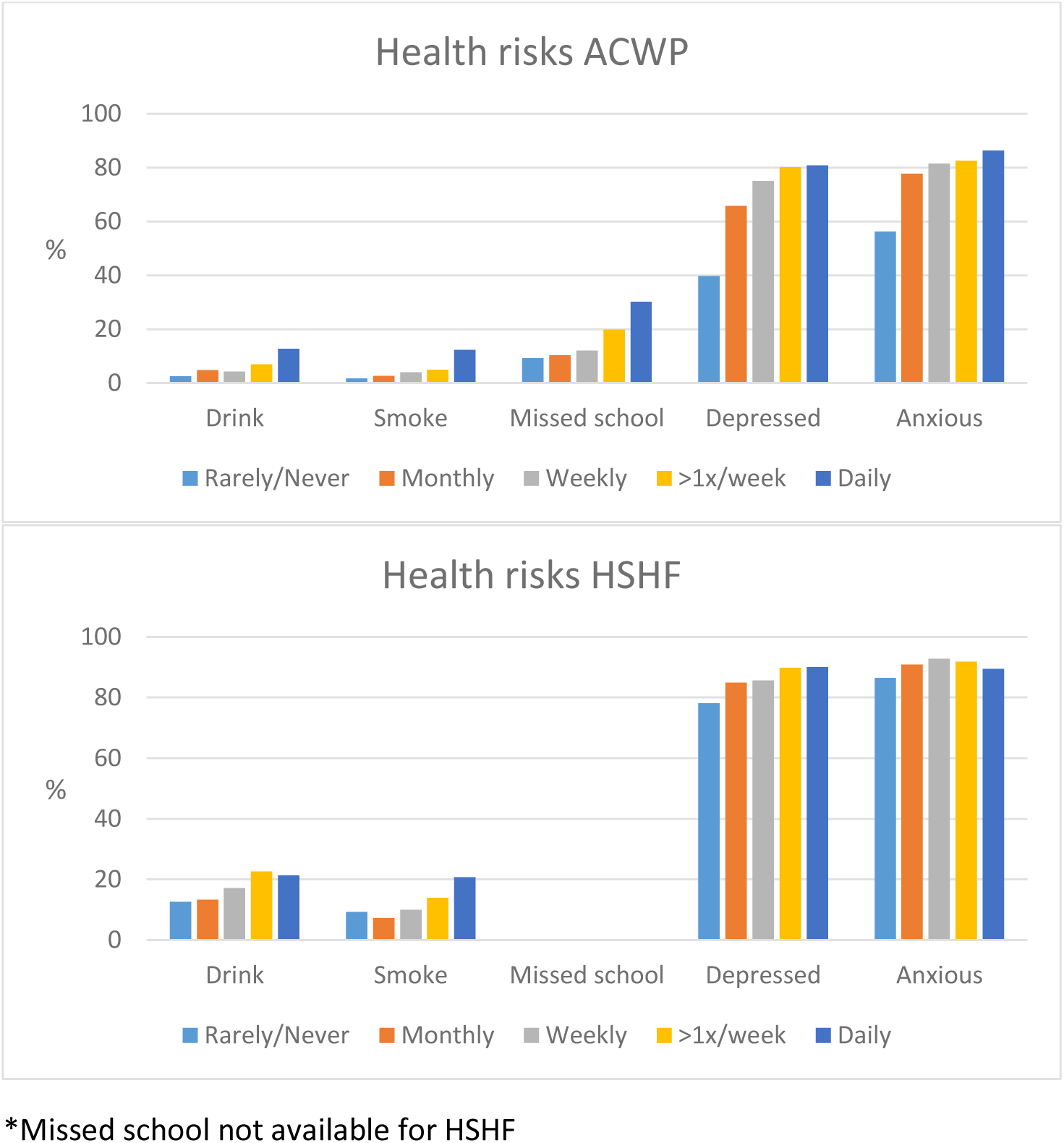
Back pain frequency and health risks

**Table 2.** Back pain frequency and health risks

| Back pain frequency | Smoking | Drinking | Depression | Anxiety | Missed school |
| --- | --- | --- | --- | --- | --- |
| **ACWP** |  |  |  |  |  |
| Rarely or never | 1.7% | 2.5% | 39.7% | 56.2% | 9.3% |
| Every month | 2.6% | 4.8% | 65.8% | 77.7% | 10.3% |
| Every week | 4.0% | 4.3% | 75.1% | 81.5% | 12.0% |
| >1x / week | 4.9% | 7.0% | 80.1% | 82.6% | 19.8% |
| Every day | 12.3% | 12.7% | 80.8% | 86.4% | 30.2% |
| * | 67.7 | 51.9 | 349.7 | 196.4 | 75.0 |
|  | p<0.01 | p<0.01 | p<0.01 | p<0.01 | p<0.01 |
| **HSHF** |  |  |  |  |  |
| Rarely or never | 9.3% | 12.6% | 78.1% | 86.5% | - |
| Every month | 7.3 % | 13.3% | 84.9% | 90.9% | - |
| Every week | 10.0% | 17.2% | 85.6% | 92.8% | - |
| >1x / week | 14.0% | 22.6% | 89.8% | 91.9% | - |
| Every day | 20.8% | 21.3% | 90.0% | 89.4% | - |
| * | 17.67 | 6.17 | 27.9 | 6.7 | - |
|  | p<0.01 | p=0.01 | p<0.01 | p<0.01 | † |

\* Mantel-Haenszel test-for-trend; † Missed school not available for HSHF

## Discussion

### Main findings

In both samples, more frequent back pain was associated with adverse health risk indicators. In most cases there was an apparent dose-response relationship as pain frequency increased, as indicated by the significant test-for-trend findings. These data indicate that back pain forms part of a wider picture of poor health risk profile. There were key differences between the datasets, in that the HSHF study preferentially recruited schools from socioeconomic strata lower than the Australian population norm, and that the mean age of the HSHF sample was approximately one year older. Although the overall rate of adverse health risk indicators and frequency of back pain report was higher in this sample, the relationship with back pain frequency was the same. This provides support for generalisability of the findings.

There are several potential explanations for the relationships observed. These include causal pathways that flow in either direction, the action of confounders or complex feedback loops between pain and the other adverse health risk indicators. The cross-sectional structure of the data does not allow us to explore the form of these relationships. However, a narrow focus on aetiology may obscure important considerations. Report of frequent pain appears to be a marker of other potentially less visible issues, such as use of alcohol and tobacco, and poor mental health. This information could be used to identify an at-risk population to target interventions. For adolescents seeking care for back pain, these health risk factors may be related to poor long-term outcome of their back complaints. While we cannot be sure that prognostic relationships are causal, this points to a promising role for addressing these issues in the context of their clinical care. Conversely, it is plausible to suggest that the presence of pain is a barrier to the effectiveness of public health interventions aimed at promoting healthy lifestyle habits, related to physical activity, substance use and mental health. This may point to a place for addressing pain in the context of broader public health interventions.

### Interpretation

These findings confirm previous literature that identifies associations between MSK pain and other indicators of poor health and health risk. As is the situation in research in adult populations, systematic reviews show a robust relationship between MSK pain and indicators of poor mental health, including depression and anxiety.^17, 22^ There is inconsistency with regard to reported relationships between adolescent pain and alcohol use and tobacco. This study provides supporting evidence that such an association is real, a conclusion that is strengthened by the dose-response relationship between frequency and likelihood of drinking or smoking apparent in both datasets.

There are numerous potential explanations for the relationships between pain and health risk factors. Tobacco use may increase pain levels via physiological pro-inflammatory and vasoconstriction processes.^7^ Alternately experience of pain may see adolescents seeking analgesic effects of substance use as has been demonstrated in adult populations.^33^ Further, behavioural or cultural influences may lead to both pain report and alcohol use or smoking, for example a study has shown maternal influences on self-medication behaviours for pain management and coping in children.^19^ Similarly, anxiety or depression could be a consequence of frequent pain experience, or make pain more likely via behavioural or cognitive processes. Socioeconomic, or genetic factors may also provide a common explanatory pathway for psychological dysfunction and pain.

### Strengths and limitations

The findings from this study come from two, relatively large and independently collected datasets. The fact that analyses reported similar relationships between the factors of interest is a strength of the study. Despite the combined sample of over 6,000 participants, data was collected from only one country, which means that findings may not generalise beyond Australia. This may be of particular issue in countries where societal attitudes and access to alcohol and tobacco use are different. The study also makes use only of cross-sectional data, this means that the findings are unsuited to drawing conclusions about causal relationships.

### Implications

These findings may have implications for clinical practice. They point to the fact that adolescents with frequent pain are at increased risk of other health problems. In the event that these other health risks influence the prognosis of painful conditions, then addressing health-related behaviours and mental health issues should form part of clinical management. Even if not causally-related to the course of MSK pain, there is an argument that comprehensive health care of the individual should include management of these factors. In either case, screening for health-related risk behaviours and indicators of poor mental health is indicated in adolescents with frequent MSK pain.

From a research perspective this study suggests that consideration of behavioural health risks, and mental health should be incorporated into research aimed at understanding the pathology of MSK pain in adolescents. Understanding these relationships will also require longitudinal studies, with careful planning of the timing of data collection. These findings also point toward the potential for developing and evaluating treatment models that integrate best-practice clinical management of pain with interventions that target lifestyle, health-related behaviours and mental health^32^

Finally, public health interventions commonly target substance use, and mental health in children and adolescents but they do not address pain. If it is the case that the same children that are at-risk of poor health outcomes due to substance use and/or mental health issues are also likely to experience pain more frequently, this may have implications for the design and content of public health interventions.

### Conclusions

Adolescents that report more frequent pain are also more likely to report higher rates of adverse health risk behaviours and indicators of poor mental health. Back pain may play a role in characterising poor overall health and risk of chronic disease in adulthood.

## References

1. Balogun O, Koyanagi A, Stickley A, Gilmour S, Shibuya K. Alcohol consumption and psychological distress in adolescents: a multi-country study. Journal of Adolescent Health. 2014;54:228–34.

2. Brattberg G. Do pain problems in young school children persist into early adulthood? A 13-year follow-up. European Journal of Pain. 2004;8:187–99.

3. Caes L, Fisher E, Clinch J, Tobias JH, Eccleston C. The role of pain-related anxiety in adolescents’ disability, and social impairment: ALSPAC data. European Journal of Pain. 2015;19:842–51.

4. Copeland WE, Shanahan L, Costello EJ, Angold A. Childhood and adolescent psychiatric disorders as predictors of young adult disorders. Arch Gen Psychiatr. 2009;66:764–72.

5. Degenhardt L, Stockings E, Patton G, Hall WD, Lynskey M. The increasing global health priority of substance use in young people. Lancet Psychiatr. 2016;3:251–64.

6. Dissing KB, Hestbaek L, Hartvigsen J, Williams CM, Kamper SJ, Boyle E, et al. Spinal pain in Danish school children – how often and how long? The CHAMPS Study-DK. BMC musculoskeletal disorders. 2017;18:67.

7. Ditre JW, Brandon TH, Zale EL, Meagher MM. Pain, Nicotine, and Smoking: Research Findings and Mechanistic Considerations. Psychol Bull. 2011;137:1065–93.

8. Gobina I, Villberg J, Villerusa A, Välimaa R, Tynjälä J, Ottova-Jordan V. Self-reported recurrent pain and medicine use behaviours among 15-year olds: results from the international study. European Journal of Pain. 2015;19:77–84.

9. Hall WD, Patton G, Stockings E, Wier M, Lynskey M, Morley KI, et al. Why young people’s substance use matters for global health. Lancet Psychiatr. 2016;3:265–79.

10. Harreby M, Kjer J, Hesselsoe G, Neergaard K. Epidemiological aspects and risk factors for low back pain in 38-year-old men and women: a 25-year prospective cohort study of 640 school children. European spine journal : official publication of the European Spine Society, the European Spinal Deformity Society, and the European Section of the Cervical Spine Research Society. 1996;5(5):312–8.

11. Heaps N, Davis MC, Smith AJ, Straker LM. Adolescent drug use, psychosocial functioning and spinal pain. J Health Psychol. 2011;16:688–98.

12. Henschke N, Harrison C, McKay D, Broderick C, Latimer J, Britt H, et al. Musculoskeletal conditions in children and adolescents managed in Australian primary care. BMC musculoskeletal disorders. 2014;15:164.

13. Hestbaek L, Leboeuf-Yde C, Kyvik KO. Are lifestyle-factors in adolescence predictors for adult low back pain? A cross-sectional and prospective study of young twins. BMC musculoskeletal disorders. 2006;7:27.

14. Hestbaek L, Leboeuf-Yde C, Kyvik KO. Is comorbidity in adolescence a predictor for adult low back pain? A prospective study of a young population. BMC musculoskeletal disorders. 2006;7:29.

15. Hodder RK, Freund M, Bowman J, Wolfenden L, Campbell E, Dray J, et al. Effectiveness of a pragmatic schoolbased universal resilience intervention in reducing tobacco, alcohol and illicit substance use in a population of adolescents: cluster-randomised controlled trial. BMJ Open. 2017;7:e016060.

16. Home Office National Statistics (UK). Drug Misuse: Findings from the 2014/2015 Crime Survey for England and Wales https://www.gov.uk/government/uploads/system/uploads/attachment_data/file/450181/drug-misuse-1415.pdf2015.2015 [

17. Huguet A, Tougas ME, Hayden J, McGrath PJ, Stinson JN, Chambers CT. A systematic review with meta-analysis of childhood and adolescent risk and prognostic factors for musculoskeletal pain. Pain. 2016;157:2640–56.

18. Institute for Health Metrics and Evaluation. GBD Compare | Viz Hub https://vizhub.healthdata.org/gbd-compare/: University of Washington; 2016

19. Jensen JF, Gottschau M, Siersma VD, Graungaard AH, Holstein BE, Knudsen LE. Association of maternal self-medication and over-the-counter analgesics for children. Pediatrics. 2014;133:e291–98.

20. Jussila L, Paananen M, Näyhä S, Taimela S, Tammelin T, Auvinen J, et al. Psychosocial and lifestyle correlates of musculoskeletal pain patterns in adolescence: A 2-year follow-up study. European Journal of Pain. 2014;18(139-46).

21. Kamper SJ, Henschke N, Hestbaek L, Dunn KM, Williams CM. Musculoskeletal pain in children and adolescents. Braz J Phys Ther. 2016;20:275–84.

22. Kamper SJ, Yamato TP, Williams CM. The prevalence, risk factors, prognosis and treatment for back pain in children and adolescents: An overview of systematic reviews. Best Prac Res Clin Rheumatol. 2016;30:1021–36.

23. Kovacs FM, Gestoso M, MT Gil del Real, López J, Mufraggi N, Méndez JI. Risk factors for nonspecific low back pain in schoolchildren and their parents: a population based study. Pain. 2003;103:259–68.

24. Law EF, Bromberg MH, Noel M, Groenewald C, Murphy LK, Palermo TM. Alcohol and tobacco use in youth with and without chronic pain. Journal of Pediatric Psychology. 2015;40:509–16.

25. Levy S, Williams JF. Adolescent substance use: the role of the medical home. Adolescent Medicine State of the Art Reviews. 2014;25:1–14.

26. Lietz P, O’Grady E, Tobin M, Murphy M, Macaskill G, Redmond G, et al. The Australian Child Wellbeing Project: Technical Report. Flinders University, the University of NSW and the Australian Council for Educational Research; 2015.

27. O’Sullivan PB, Beales DJ, Smith AJ, Straker LM. Low back pain in 17 year olds has substantial impact and represents an important public health disorder: a cross-sectional study. BMC public health. 2012;12:100.

28. Shiri R, Karppinen J, Leino-Arjas P, Solovieva S, Viikari-Juntura E. The association between smoking and low back pain: a meta-analysis. American Journal of Medicine. 2010;87:e7–35.

29. Swain MS, Henschke N, Kamper SJ, Gobina I, Ottová-Jordan V, Maher CG. An international survey of pain in adolescents. BMC public health. 2014;14:447.

30. Taylor M, Collin SM, Munafò MR, MacLeod J, Hickman M, Heron J. Patterns of cannabis use during adolescence and their association with harmful substance use behaviour: findings from a UK birth cohort. J Epidemiol Community Health,. 2017;71:764–70.

31. Weissman DG, Schriber RA, Fassbender C. Earlier adolescent substance use onset predicts stronger connectivity between reward and cognitive control brain networks. Dev Cogn Neurosci. 2015;16:121–29.

32. Williams A, Wiggers J, O’Brien KM, Wolfenden L, Yoong S, Campbell E, et al. A randomised controlled trial of a lifestyle behavioural intervention for patients with low back pain, who are overweight or obese: study protocol. BMC musculoskeletal disorders. 2016;17:70.

33. Zale EL, Maisto SA, Ditre JW. Interrelations between pain and alcohol: an integrative review. Clin Psychol Rev. 2015;37:57–71.

